# Genomic diversity of *Streptococcus uberis* isolated from clinical mastitis of cattle in selected areas of Bangladesh

**DOI:** 10.1101/2022.12.19.521080

**Authors:** Jayedul Hassan, Md. Abdus Sattar Bag, Ajran Kabir, Md. Wohab Ali, Maqsud Hossain, Md. Tanvir Rahman, Md. Shafiqul Islam, Md. Shahidur Rahman Khan

## Abstract

Streptococci are the major etiology in mastitis, a cause of huge economic losses in the dairy industries. *Streptococcus* (*S*.) *agalactiae, S. dysagalactiae* and *S. uberis* are mostly encountered in bovine mastitis; however, data on the diversity and characteristics of Streptococcus in clinical mastitis of cattle in Bangladesh is lacking. Thus, the present study was aimed to determine the diversity and antimicrobial resistance pattern of *Streptococcus* spp. isolated from clinical mastitis of cattle reared in Bangladesh. A total of 105 milk samples comprising eighty (80) from cattle with clinical mastitis (CCM) and twenty-five (25) from apparently healthy cattle (AHC) in four prominent dairy farms and one dairy community were purposively collected and examined in this study. Milk samples were enriched in Luria Bertani broth (LB) and *Streptococcus* spp. was isolated on Modified Edwards Medium and identified by 16S rRNA gene sequencing. Among eighty (80) clinical samples, eighteen (18) were positive for *Streptococcus* spp. while none of the milk from AHC revealed Streptococcus by cultural and molecular examination. Sequencing and phylogenetic analysis identified 55.6%, 33.3%, 5.6% and 5.6% of the Streptococcus isolates as *S. uberis, S. agalactiae, S. hyovaginalis* and *S. urinalis*, respectively. Antibiotic sensitivity testing with antimicrobials commonly used to treat clinical mastitis revealed 100%, 100%, and 30% of the *S. agalactiae*, *S. hyovaginalis*, and *S. uberis* as multidrug-resistant, respectively. Molecular characterization through whole genome sequencing of five (5) *S. uberis* isolates identified at least two novel ST types of *S. uberis* circulating in the study areas with one ST (4/5 isolates) clustered with the isolates from China, India and Thailand, and the other (1/5) with UK, Ireland and Australia. Pan-genome analysis and phylogeny of the core genome sequences also clustered the isolates into two sub-clusters, indicating the presence of at least two different subtypes of *S. uberis* in the study area. On virulence profiling, all the isolates of this study were found to harbor at least 35 virulence and putative virulence genes probably associated with intramammary infection (IMI) indicating all the *S. uberis* isolated in this study as potential pathogen. From the overall findings it was evident that Strepococcus occurring in bovine mastitis are diverse and *S. uberis* genome carries an array of putative virulence factors which need to be investigated genotypically and phenotypically to identify a specific trait or determinant governing the virulence and fitness of this bacterium. Moreover, Streptococcus isolated in this study carried multidrug resistance which needs careful consideration during the selection of a treatment regimen for mastitis.

## INTRODUCTION

Bovine mastitis is the major hindering factor in the dairy industry responsible for huge economic losses worldwide. Mastitis was estimated to account for 125 billion dollars in losses worldwide through the reduction of milk yield and quality, culling cows and veterinary care [1]. Bacteria are responsible for the clinical onset of this disease among 130 well-known pathogens of bovine mastitis [2][3][4]. Genus Streptococcus is major among the bacterial pathogens and responsible for 23-50% of the total mastitis cases worldwide [5]. Mastitis is classified into three distinct forms, sub-clinical, clinical and chronic [1], and the occurrence of Streptococcus have been reported in all three forms of this disease [6][7][8].

*St. agalactiae, S. dysagalactiae*, and *S. uberis* are most commonly found in bovine mastitis with varying global frequencies among *Streptococcus* spp. [5][9][10]. *S. agalactiae* was found in 90% of mastitis cases in the past; however, due to their contagious nature, the incidence of *S. agalactiae* was drastically reduced by implementing a five-point hygiene plan in the 1960s [2][11] [12]. In contrast, the majority of the mastitis cases shifted towards environmental mastitis [2][13][14]. As a result, the prevalence of environmental mastitis pathogens such as *S. uberis* has increased remarkably [5][15][16]. However, the prevalence of the major streptococci was variable based on geographical locations [5]. *S. uberis* is most widespread in Australia, New Zealand, and North America, while *S. agalactiae* was ubiquitous in Africa and Asia. In South Africa, the incidences of both pathogens are comparable [5]. The difference in prevalence might largely depend on environmental factors and farm-specific practices, including local outbreaks [5]. In Bangladesh, information on Streptococcus and their antibiotic susceptibility in mastitis are minimal [17][18][19], and to the best of our knowledge, no research has been documented on the occurrence, diversity and molecular characteristics of Streptococcus in clinical mastitis of cattle in Bangladesh. Thus, the present study aimed to determine the diversity of *Streptococcus* spp. in clinical mastitis of cattle in the major dairy production units (farms/ community) in Bangladesh and their antibiotic susceptibility pattern. In addition, the research was further extended to the genomic characterization and virulence profiling of the *S. uberis* isolated in this study through whole genome sequencing and subsequent bioinformatic analysis considering the increased prevalence of this pathogen in bovine mastitis.

## Materials and methods

### Sample collection and enrichment

Milk samples were collected from cattle with clinical mastitis (CCM) previously diagnosed by the field or farm veterinarian from four major dairy farms of Dhaka and Mymensingh district and one dairy community in Sirajganj district of Bangladesh [20]. Through a single visit per farm milk samples from all the CCM were directly collected from the udder aided by the resident veterinarian. In addition, milk samples from cattle with no visible sign of mastitis or negative to California Mastitis test (CMT) (herein stated as apparently healthy cattle (AHC)) were also examined to ascertain the variable characteristics of *Streptococcus* spp. in AHC and CCM. A total of 105 milk samples were examined in this study, comprising 80 from CCM and 25 from AHC, respectively. A 10 ml of composite milk sample was collected directly from the udder by the residential veterinarian of the respective farm and carried to the laboratory in an ice box for microbiological analysis. In the laboratory, the samples were enriched in LB broth as described previously [21].

### Isolation and identification of *Streptococcus* spp

One hundred microlitres (100 μl) of enriched culture was spread onto modified Edwards medium (MEM) (Himedia, India) and incubated overnight at 37°C [20]. Colonies characteristic to *Streptococcus* spp. (blue and/ or black color) were screened by Gram’s staining followed by purification to single isolated colonies by subsequent streaking onto MEM. Crude DNA was extracted from the isolated colonies using the boiling method and confirmed as Streptococcus by PCR and sequencing of 16S rRNA using 8F (5’-AGAGTTTGATCMTGGC-3’) and 1492R (5’-TACCTTGTTACGACTT-3’) primers [22]. Alignment and phylogenetic analysis of the 16S rRNA sequences were performed on CLC genomic workbench 22 (Qiagen, Germany).

### Antibiotic susceptibility testing

Susceptibility of Streptococcus against antibiotics commonly used in mastitis treatment was performed by the disc diffusion method [23]. A total of 14 antimicrobials of different classes including β-lactams (amoxicillin 10 μg (AMX), ampicillin 10 μg (AMP), Ceftazidime 30 μg (CAZ), Ceftriaxone 30 μg (CTR), Cephalexin 30 μg (CN), and penicillin G 10 μg (P)), aminoglycosides (kanamycin 30 μg (K), gentamicin 30 μg (GEN), and streptomycin 10 μg (S)), glycopeptides (vancomycin 30 μg (VA)), macrolides (azithromycin 15 μg (AZM)), phenicols (chloramphenicol 30 μg (C)), polypeptides (bacitracin 10 units (B)), and tetracyclines (tetracycline 30 μg (TE)) was used in this study. *E. coli* strain ATCC25922 was used as the control strain and the results were interpreted according to the Clinical and Laboratory Standards Institute (CLSI) guidelines [24]. Isolates exhibiting resistance to three or more antibiotic classes were considered as multidrug-resistant (MDR) [25].

### Whole genome sequencing and annotation

Five of ten *S. uberis* isolated in this study were subjected to whole genome sequencing (WGS). The isolates were selected covering all the study areas to have a diverse picture of the genomes. WGS was performed on an Illumina Nextseq 550 platform (Illumina, CA, USA). Quality of the FASTQ reads and trimming of low-quality residues was performed on Trimmomatic (Galaxy Version 0.38). Trimmed reads were assembled on a hybrid assembler Unicycler (Galaxy Version 0.4.8.0). Assembled contigs were annotated using Prokka (Galaxy Version 1.14.6), RAST (Rapid Annotation using Subsystem Technology) (https://rast.nmpdr.org/rast.cgi) and NCBI prokaryotic genome annotation pipeline (PGAP) to identify the functional features. The annotated data were used for antibiotic resistance gene profiling, plasmid identification, sequence typing and virulence genes profiling,

Antibiotic resistance genes were identified using the Comprehensive Antibiotic Resistance Database (CARD) (https://card.mcmaster.ca/), and for virulence gene detection, the *S. uberis* Putative Virulence Database suPVDB [26] virulence factor database (VFDB) (www.mgc.ac.cn/VFs) was explored. Further, the known virulence related genes [27][28][29]; were identified through search on NCBI blastn (Nucleotide BLAST: Search nucleotide databases using a nucleotide query (nih.gov) and blastx (blastx: search protein databases using a translated nucleotide query (nih.gov)) platforms.

Average nucleotide identity (ANI) of the *S. uberis* sequences along with genome assemblies downloaded from the NCBI (Streptococcus uberis - Assembly - NCBI (nih.gov)) was calculated using a perl program GET_HOMOLOGUES [30]. The genome sequences were further analyzed using the Roary Pan Genome pipeline [31] to identify orthologous genes in *S. uberis* and construction of a tree by clustering the core and accessory genes in a matrix. In addition, core genome based whole genome phylogeny and tree visualization were performed on CLC Genomic workbench (Qiagen) and iTOL v6 (iTOL: Interactive Tree Of Life (embl.de)), respectively. CRISPRCasFinder 4.2.20 software at Proksee tools (https://Proksee.ca); was used for rapid identification and classification of clustered regularly interspaced short palindromic Repeats (CRISPRs) and the PHASTER web server (https://phaster.ca) was used to identify a prophage sequence. The *S. uberis* genome sequences were examined for their sequence type (ST) through PubMLST website (pubmlst.org). Concatenated ST sequences were downloaded from PubMLST (https://pubmlst.org/organisms/streptococcus-uberis) and the minimum spanning tree was constructed using the goeBurst algorithm of Phyloviz ([PHYLOViZ Online](https://online.phyloviz.net/index))) to identify the closest neighbors.

## Results

### Occurrence and diversity of *Streptococcus* spp

*Streptococcus* spp. was detected in 18 of 105 milk samples, including 18/80 (22.5%) from CCM and 0/25 (0.0%) from AHC (Table 1). *St agalactiae, S. uberis, S. hyovaginalis*, and *S. urinalis* (Fig 1) have been identified using sequencing, homology search, and phylogenetic analysis (Accession no. OL581668 to OL581684, and ON935775). *S. uberis* was most prevalent (55.6%), followed by *S. agalactiae* (33.3%) (Table 1). None of the samples included *S. dysgalactiae*.

**Table 1.** Prevalence and antibiotic resistance pattern of the *Streptococcus* spp. isolated in this study.

| Species | Strain | Sample positive (%)<br>(n = 18) | Antibiotic sensitivity pattern | MDR |
| --- | --- | --- | --- | --- |
| *S. agalactiae* | 2010 | 33.33 | AZM-CAZ-CIP-CTR-GEN-K-P-S-TET | 55.6% |
|  | 2013 |  | CIP-K-P |  |
|  | 2014 |  | CAZ-CIP-CN-CTR-GEN-K-P-S-TET |  |
|  | 2087 |  | CAZ-CIP-CTR-GEN-K-P-S-TET |  |
|  | 2090 |  | AMP-AMX-CIP-GEN-P-TET |  |
|  | 2095 |  | AMP-AMX-AZM-CAZ-CTR-B-P-TET |  |
| *S. uberis* | 2002 | 55.56 | AZM-CIP-CN-K-P-TET |  |
|  | 2021 |  | CIP-CN-CTR-S-TET |  |
|  | 2053 |  | CAZ-CIP-CTR-S |  |
|  | 2055 |  | CIP-TET |  |
|  | 2058 |  | CIP-TET |  |
|  | 2060 |  | P-TET |  |
|  | 2062 |  | CIP-TET |  |
|  | 2063 |  | CIP-TET |  |
|  | 2065 |  | CIP-TET |  |
|  | 2066 |  | CIP-TET |  |
| *S. hyovaginalis* | 2022 | 5.56 | CAZ-CIP-CN-CTR-GEN-K-P-S-TET |  |
| *S. urinalis* | 2042 | 5.56 | - |  |

Legends: AZM, Azithromycin; AMP, Ampicillin; AMX, Amoxicillin; B, Bacitracin; CAZ, Ceftazidime; CIP, Ciprofloxacin; CN, Cephalexin; CTR, Ceftriaxone; K, Kanamycin; GEN, Gentamicin; P, Penicillin; S, Streptomycin; TET, Tetracycline

**Fig 1.**
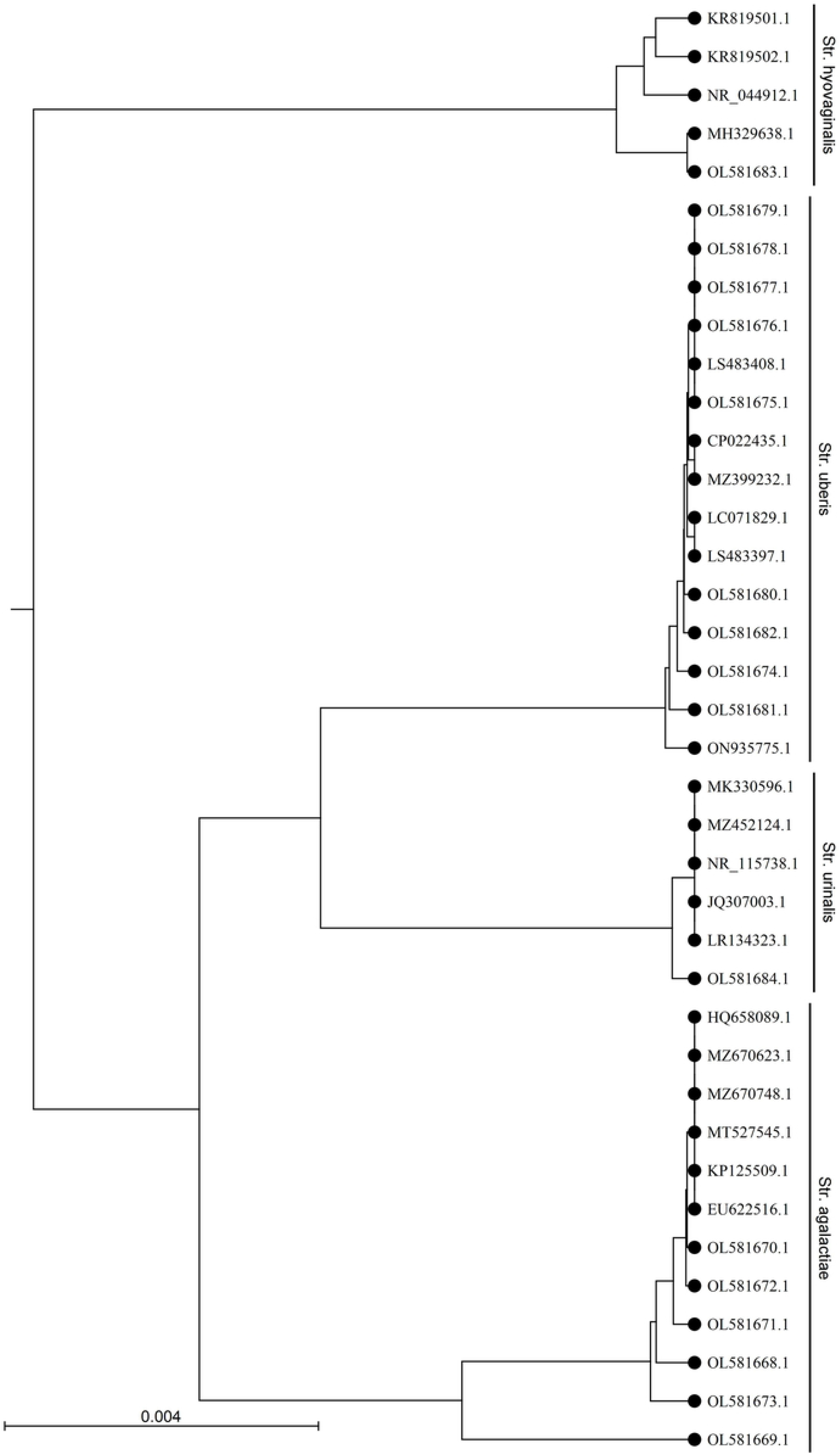
Phylogenetic analysis of the 16S rRNA gene sequences of *Streptococcus* sp. isolated in this study. The UPGMA tree was prepared using CLC sequence viewer 8.0 and the nucleotide distances were calculated using Kimura 80 model. Accession numbers with underlines denotes sequences obtained in this study.

### Antibiotic susceptibility of Streptococcus isolates

Fifty-five percent (55.6%, 10/18) of the Streptococcus isolated in this study showed multidrug resistance (MDR), with 100% of the *S. agalactiae* and *S. hyovaginalis*. Whereas, *S. urinalis* was susceptible to all the antimicrobials used in this study (Table 1). On the other hand, only 30% of the *S. uberis* showed MDR, and they were susceptible to most of the antibiotics used in this study (Table 1). Among the antibiotics used, the highest resistance was encountered against ciprofloxacin (83.3%) and tetracycline (83.3%). All (100%) of the Streptococcus isolates were sensitive to chloramphenicol and vancomycin (Table 1).

### General features of the *S. uberis* genomes

Five *S. uberis* isolated in this study (BD isolates) were sequenced and assembled to elucidate their genomic features. The assembled genome length ranged from 18,86,469 to 20,33,655 bp with GC contents of 36.4 to 36.5% (Table 2). The draft genome of strain 2053 was the largest and strain 2055 being the smallest among the genomes examined in this study. The genomes consisted of 1840 to 2032 coding sequences (CDS) and 44 to 45 RNAs. Strains 2002, 2021, 2055 and 2062 carried 100% identical nucleotide sequences, but 2053 shared 99.04% sequence identity with them (Fig 2). The isolates carried 1628 core genes and 645 shell genes (Fig 3a). Gene clustering based on the presence and absence of genes in *S. uberis* revealed identical patterns of the genome 2002, 2021, 2055 and 2062 on a pangenome matrix (Fig 3b) and clustered close to the pathogenic strain 0140J, whereas the strain 2053 clustered close to the non-pathogenic strain EF20 described from the UK (Hossain et al., 2015) (Fig 3b). Similar pattern of clustering was evident on a phylogenetic tree constructed with core genome sequences of *S. uberis* genomes described throughout the world (Fig 3c).

**Table 2.**
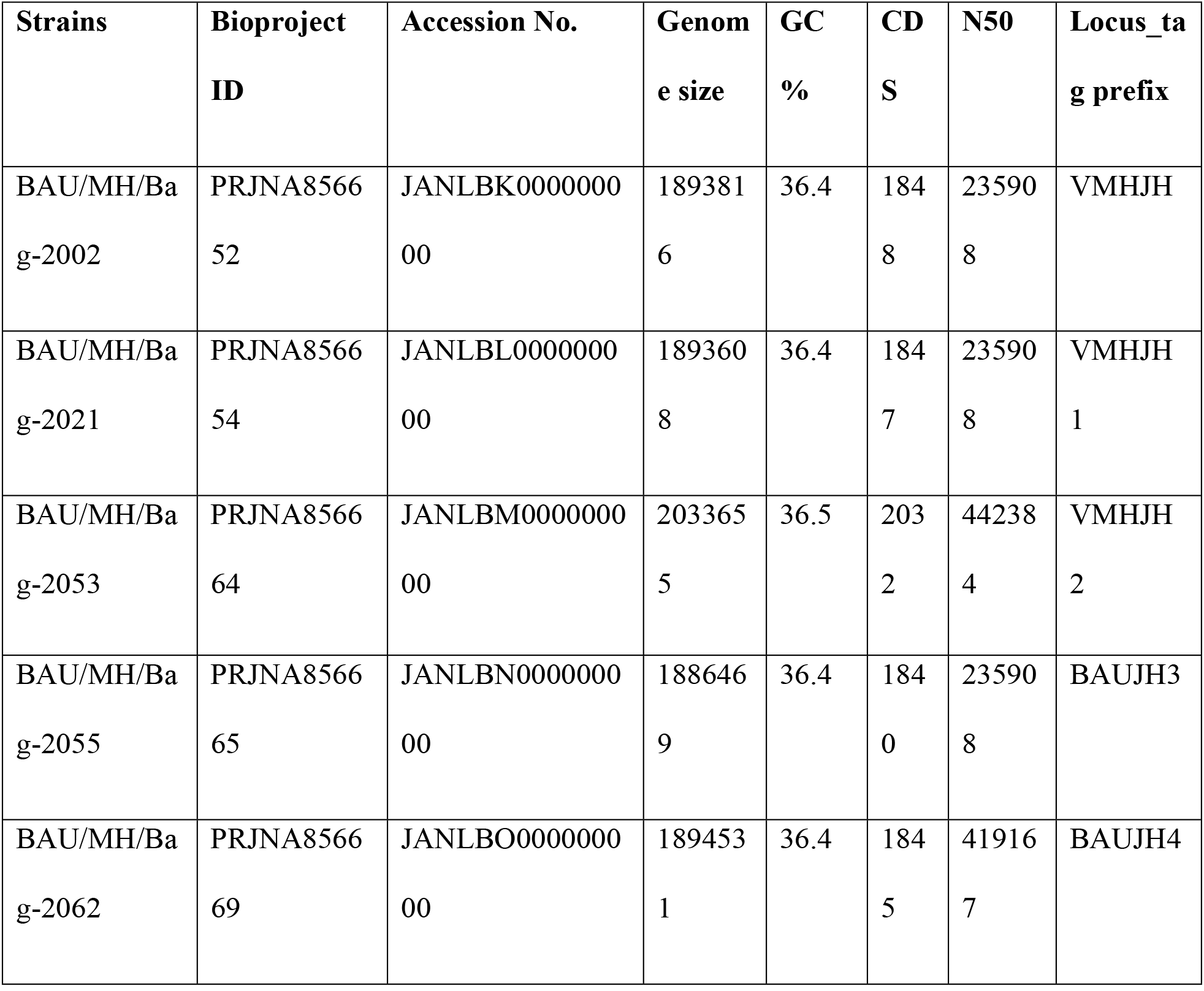
WGS information of the *Streptococcus uberis* isolated in this study.

**Fig 2.**
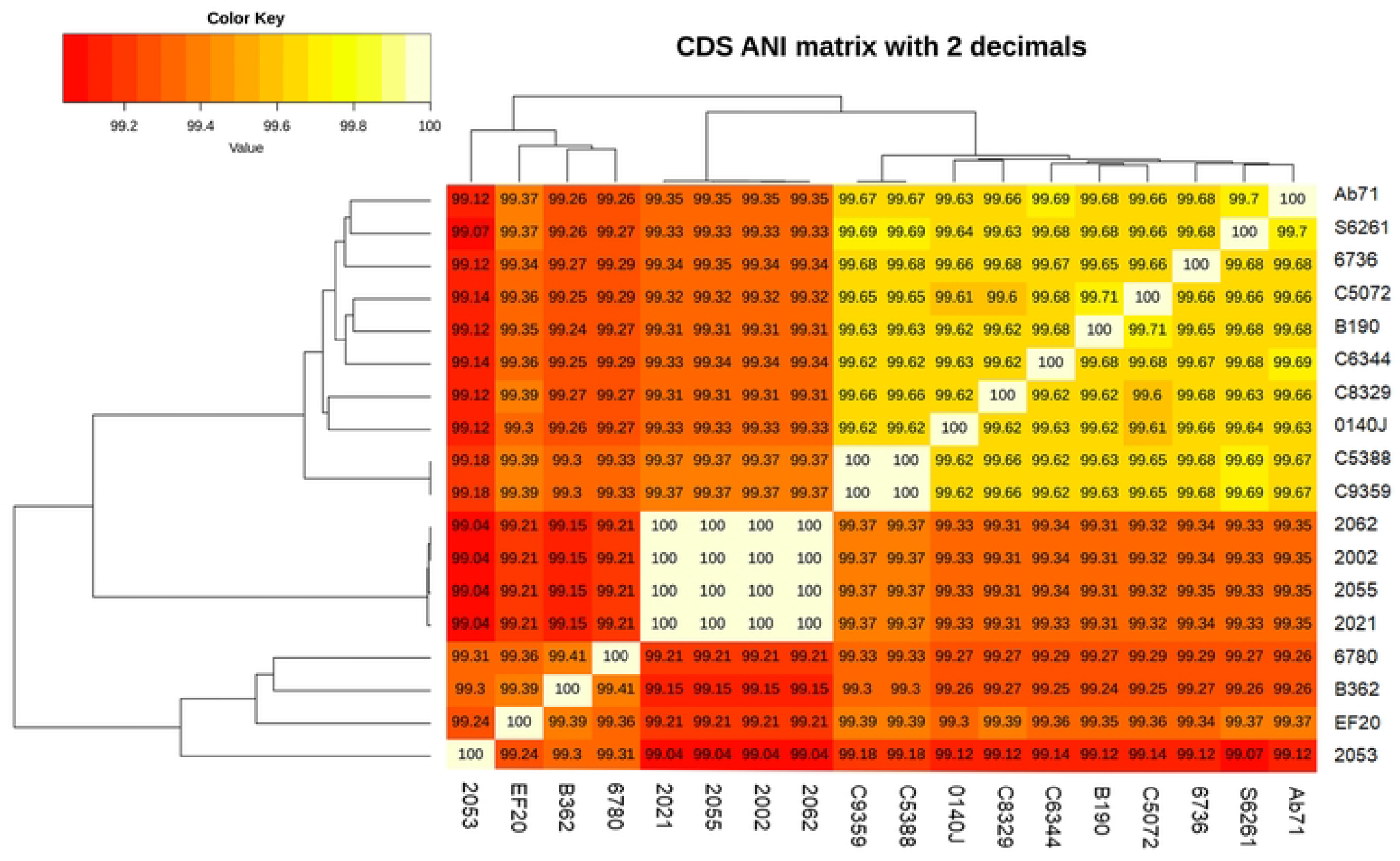
Average nucleotide identity (ANI) of *Streptococcus uberis* genomes described in this study and related strains from UK (Hossain et al., 2015). The nucleotide identity was calculated with get_homologue.pl.

**Fig 3.**
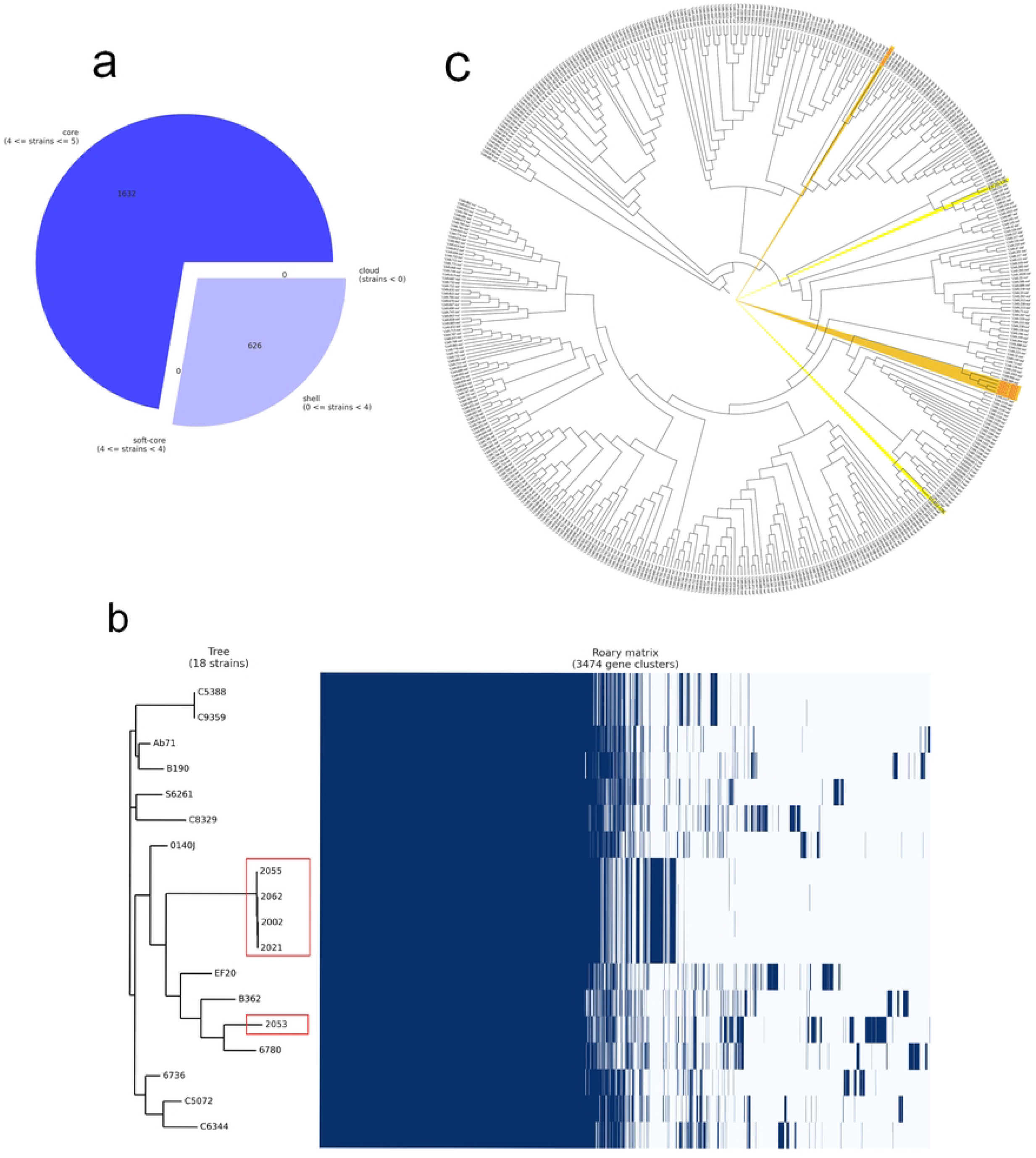
Pangenome analysis of the *Streptococcus uberis* isolated from Bangladesh (BD). a) breakdown of genes in *S. uberis* isolates from BD, b) pangenome based (gene presence and absence) gene clustering matrix of BD (enclosed in red boxes) and UK isolates and c) core genome based phylogeny with *S. uberis* genomes described throughout the world. The figures were prepared based on the data acquired from Roary Pangenome analysis using roary_plots.py script.

### Sequence type (ST) of the *S. uberis* isolates

Whole genome based sequence typing through the PubMLST website revealed the existence of two novel ST sequences in the *S. uberis* isolated from BD. One ST is close to ST474 (isolates 2002, 2021, 2055, and 2063), and the other (2053) was close to several ST sequences of ST152, ST133, ST82 and ST968. Phylogeny of the concatenated ST sequences clustered the former group with sequences described from India, Thailand and China. On the other hand, ST sequences of 2053 closely clustered with isolates from Ireland, UK and Australia (Fig 4).

**Fig 4.**
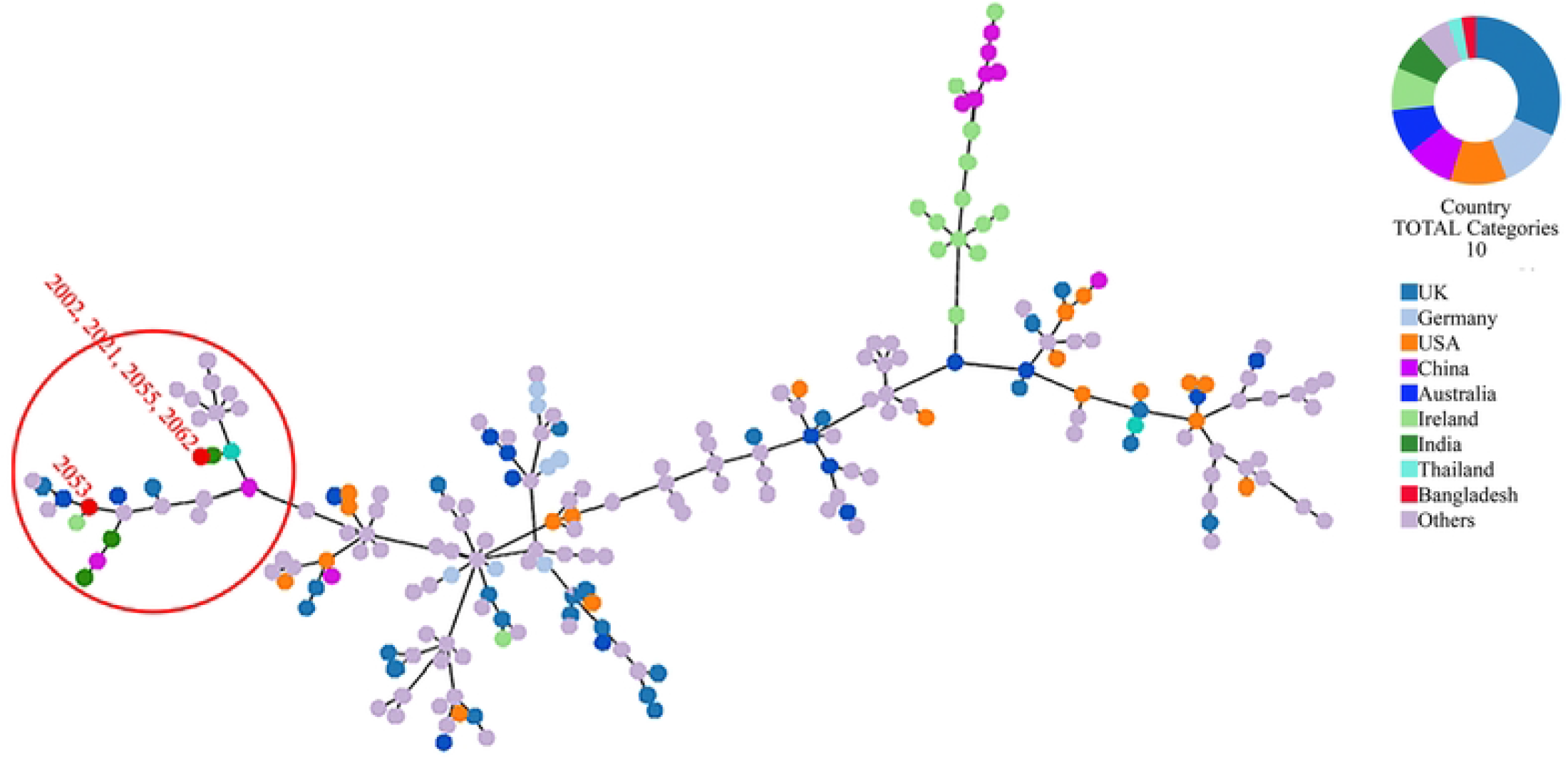
MLST phylogeny of *S. uberis* sequences obtained in this study. Concatenated ST sequences of *S. uberis* were downloaded from PubMLST (https://pubmlst.org/organisms/streptococcus-uberis) and the minimum spanning tree was constructed using goeBurst algorithm of Phyloviz ([PHYLOViZ Online](https://online.phyloviz.net/index)). Red dot inside the circle denotes isolates described in this study from Bangladesh.

### Virulence factor and antibiotic resistance in the *S. uberis* isolates

The *S. uberis* genomes were examined for the presence of sixty-one (61) putative virulence genes described so far. The virulence factors and corresponding orthologs identified in the *S. uberis* genomes are provided in Fig 5 and S1 Table. All the BD isolates carried thirty-five (35) of the virulence genes examined. In addition, strains 2002, 2021, 2055 and 2062 carried five virulence genes such as *cps4E/pglC, cpsM,, hasA, hasB*, and uberolysin encoding genes which were absent in strain 2053. On the other hand, 2053 carried one putative virulence factor, “collagen-like surface-anchored protein”, which was absent in the 4 other isolates. Analysis of the *has* operon revealed that all four positive isolates carried *hasA* and *hasB* gene in chronicles, but the *hasC* gene was located 3.9 kb downstream of the *hasAB* and in the opposite direction. In addition, upstream of the VMHJH_06810, VMHJH1_07130, VMHJH2_06655, BAUJH3_05795, BAUJH4_06250 locus of the BD isolates, which were homologous to *vru* (SUB0144) of strain 0140J was analyzed for a four bp deletion region as described by Hossain et al., 2015. Unlikely 0140J, a five bp (TATAA) deletion was observed in the position −75 to −79 in all the *S. uberis* isolates described in this study (Fig 6).

**Fig 5.**
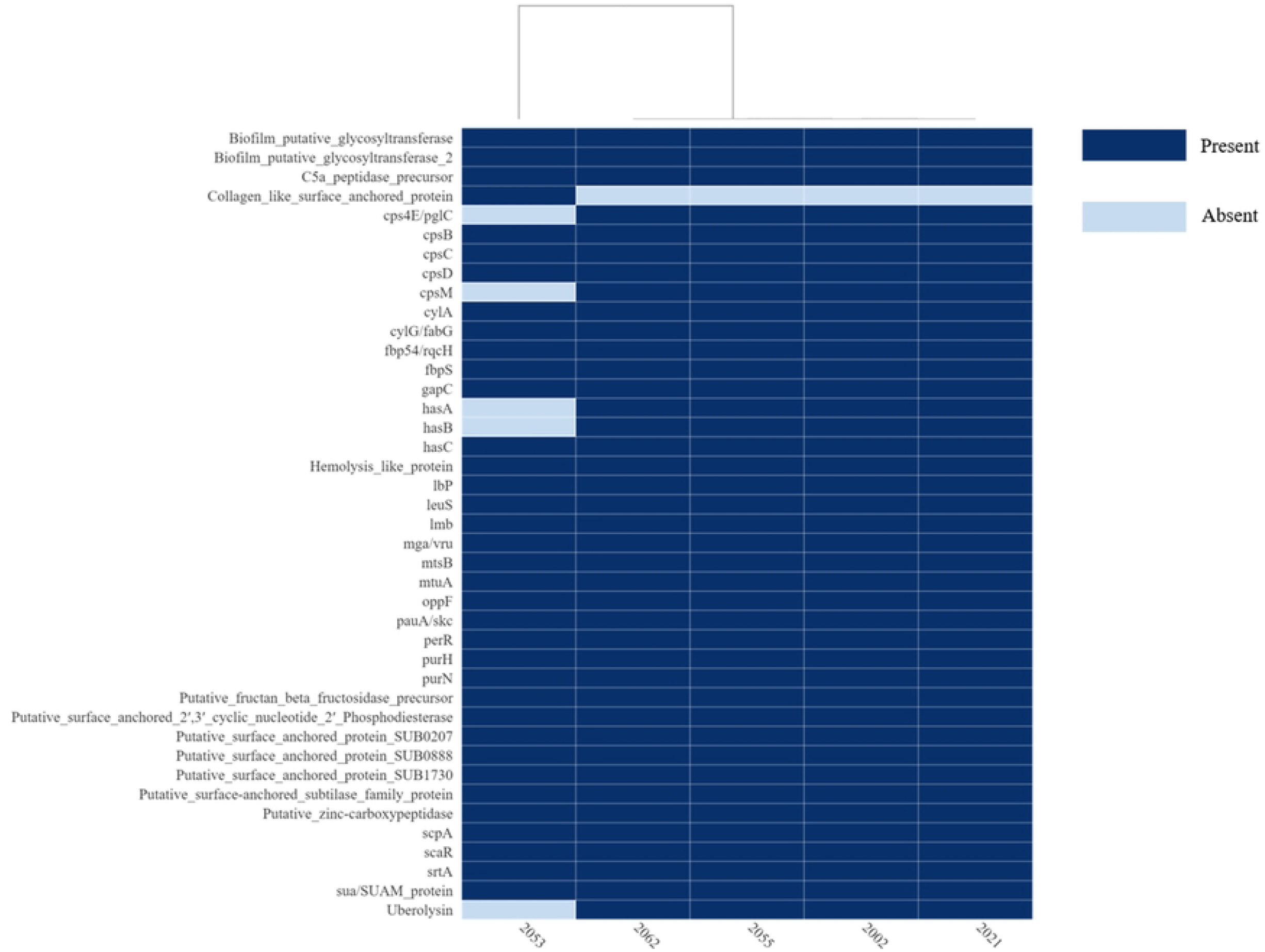
Heat map of the virulence or putative virulence genes/ factors present or absent in the *Streptococcus uberis* genomes from Bangladesh. The isolate ID is in the x-axis and the virulence genes/ factors are in the y-axis. The figure was prepared on DISPLAYR (https://southeastasia.displayr.com/) using default parameters and dendrogram appearance.

**Fig 6.**
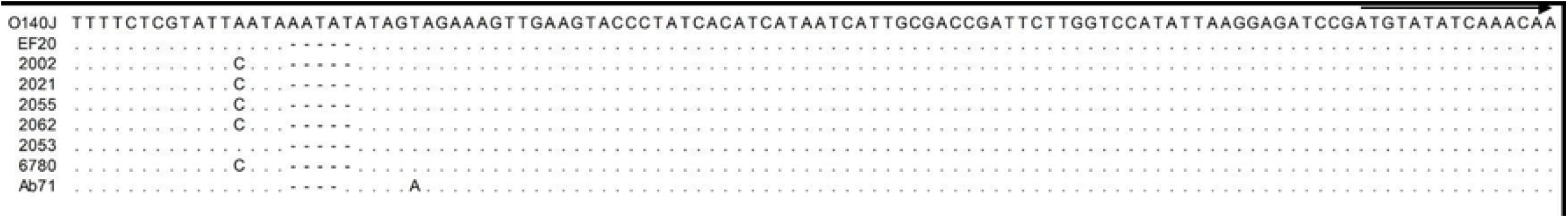
Analysis of the upstream of *vru* gene homologues in the *Streptococcus uberis* isolated in this study. Upstream of the *vru* was identified according to the criteria described by Hossain et al., 2015 and the alignment was performed on CLC Genomic Workbench 22.0. The initiation of the vru homologue is indicated with arrow. Matching residues were replaced with dots and the deleted residues are marked with dashes. Deletion of 5 pb TATAA was observed in all the strain (2002, 2021, 2053, 2055, 2062) described in this study.

Analysis of the *S. uberis* genome through CARD revealed the presence of *tetL, tetM, patA, patB* and *lnuD* genes in all but in strain 2053 only the *patA* and *patB* genes were evident. The genes *tetL* and *tetM* confer resistance to tetracyclines, *patA* and *patB* to fluoroquinolones and *lnuD* to lincosamide antibiotics. The antibiotic resistance genotype correlates with the phenotypic ciprofloxacin and tetracycline resistance pattern of the BD isolates (Table 1).

### Analysis of the CRISPR-Cas regions

*S. uberis* isolated in this study were examined for CRISPR-Cas regions where the region was found deleted in all the strains except 2053 when compared to that with reference strain 0140J. In strain 2053 a Type II CRISPR/Cas system was identified comprising *cas9_cas1 _cas2 _csn2-st* genetic arrangement (Fig 7).

**Fig 7.**
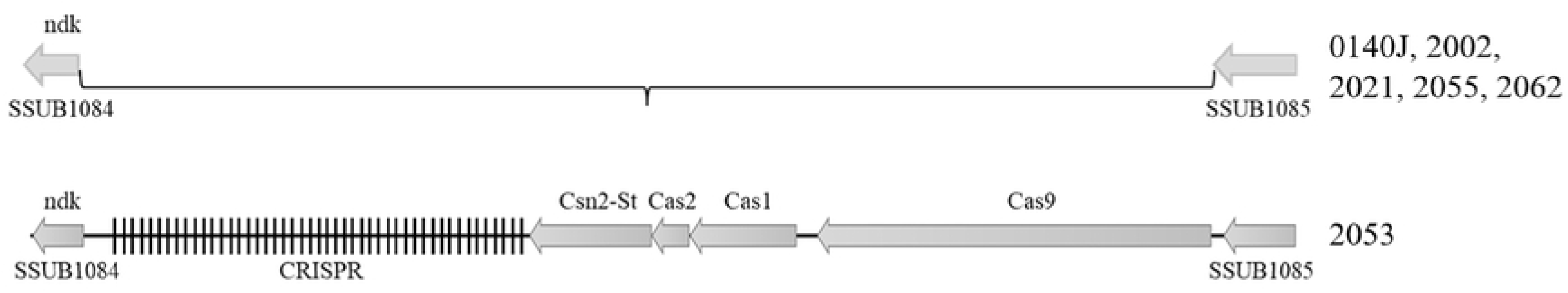
Type-II CRISPR region in the *S. uberis* isolates. Figure showing the relative position of the CRISPR-Cas genes in the *S. uberis* genome. *S. uberis* strain 0140J was used as the reference genome. The CRISPR-Cas system was detected only in strain 2053 among the 5 BD isolates.

### Prophage and other mobile genetic elements

Prophages are significant in the evolution of bacteria and their virulence. The *S. uberis* genomes were examined for the presence of prophages through the PHASTER website. All the isolates carried two to four prophages; however, intact prophages were detected only in strain 2053 (Table 3). The strain 2053 carried two intact prophages of 44.8 kb (locus_tag: VMHJH2_06750 – 07050) and 33.7 kb (locus_tag: VMHJH2_09380 – 09605) sizes with 39.17 and 37.43% GC contents, respectively. Prophages of 8.6 to 11Kb sizes were present in the other 4 isolates, but they were incomplete (Table 3). In further analysis, the intact phages contained fifty-seven and forty-four protein sequences, respectively, where most of the proteins were phage-related (forty seven and thirty five sequences, respectively). No known virulence related genes were found in the prophage sequences. The *S. uberis* strains did not carry any plasmids but carried a number of insertion sequences (Table 4). None of the insertion sequences was also associated with virulence determinants.

**Table 3.** Distribution of prophages in the *Streptococcus uberis* isolated in this study

| **Strain** | **Region** | **Length** | **Completeness** | **CDS** | **Possible phage** | **GC%** |
| --- | --- | --- | --- | --- | --- | --- |
| 2002 | 1 | 8.6Kb | Incomplete | 11 | PHAGE_Strept_9874_NC_031023 | 37.88 |
|  | 2 | 11Kb | Incomplete | 10 | PHAGE_Clostr_phiCT19406C_NC_029006 | 31.30 |
| 2021 | 1 | 8.6Kb | Incomplete | 11 | PHAGE_Escher_RCS47_NC_042128 | 37.88 |
|  | 2 | 11Kb | Incomplete | 10 | PHAGE_Bacill_CampHawk_NC_022761 | 31.30 |
| 2053 | 1 | 26.8 | Incomplete | 36 | PHAGE_Strept_SpSL1_NC_027396 | 35.25 |
|  | 2 | 20.5 | Incomplete | 22 | PHAGE_Pectob_phiA41_NC_048659 | 36.97 |
|  | 3 | 44.8 | Intact | 57 | PHAGE_Strept_A25_NC_02869 | 39.17 |
|  | 4 | 33.7 | Intact | 44 | PHAGE_Strept_2972_NC_007019 | 37.43 |
| 2055 | 1 | 8.6kb | Incomplete | 11 | PHAGE_Cronob_vB_CsaM_GAP32_NC_019401 | 37.88 |
|  | 2 | 11Kb | Incomplete | 10 | PHAGE_Rhizob_RR1_A_NC_021560 | 31.30 |
| 2062 | 1 | 8.6Kb | Incomplete | 11 | PHAGE_Escher_RCS47_NC_042128 | 37.88 |
|  | 2 | 11Kb | Incomplete | 10 | PHAGE_Bacill_PfEFR_5_NC_031055 | 31.30 |

**Table 4.** Transposon and insertion sequences in the *Streptococcus uberis* isolated in this study

| IS family | Strain ID (no. of sites) |
| --- | --- |
| IS256 | 2002, 2012, 2055, 2066 |
| IS3 | 2002 (2), 2021, 2053, 2055, 2062 |
| IS6 | 2002 (3), 2021 (3), 2055, 2062 (2) |
| unknown | 2002, 2021 (2), 2053 (2), 2055 (2), 2062 (2) |

## Discussion

The present study aimed to determine *Streptococcus* spp. in clinical mastitis of cattle in Bangladesh. Milk samples from clinical mastitis (CM) and apparently healthy cattle (AHC) were collected from major dairy farms in Dhaka, Mymensingh and Sirajganj district of Bangladesh. On cultural examination and sequencing of 16Sr RNA gene, milk from 22.5% of the CCM but none of the AHC were found positive for *Streptococcus* spp. The occurrence of Streptococcus recorded in this study is in consent with the worldwide prevalence of *Streptococcus* spp. in mastitis [5]. In this study, four different species of Streptococcus, including *S. agalactiae, S. uberis, S. hyovaginalis* and *S. urinalis* with the highest prevalence of *S. uberis* (55.56%) followed by *S. agalactiae* (33.33%) was identified. The increased occurrence of *S. uberis* is in consent with current research throughout the world [32][33][34][35]; but differs with that reported from Asia where *S. agalactiae* was predominantly associated with mastitis [5]. The difference in the occurrence of *S. uberis* in mastitis is not impossible, and to further reveal the actual scenario, a country-wide study with more samples is necessary. Besides, we are reporting two new species *S. hyovaginalis* and *S. urinalis* from clinical mastitis for the first time in the world. Although the pathogenicity or association of these two species with mastitis was not studied, they were the sole *Streptococcus* spp. isolated from the respective milk samples, indicating their possible association with mastitis; however, further studies are necessary to elucidate their relationship and virulence potential to be a mastitis pathogen.

Antimicrobial resistance (AMR) is a critical problem making mastitis control challenging through systemic or intramammary therapies. Moreover, extensive use of antimicrobials in mastitis control resulted in antimicrobial residue in milk and entering the human body through the food chain [13][36]. Mastitis caused by *Streptococcus* spp. are difficult to treat due to their ability to form abscess and survive intracellular [37][13]. However, antibiotic resistance of *Streptococcus* spp. depends on the species being studied. In this study, we found that 100% of the *S. agalactiae* and *S. hyovaginlais* were multidrug-resistant, whereas antibiotic resistance was observed in 30% of the *S. uberis* and none of the *S. urinalis*. The highest resistance was observed against Ciprofloxacin and tetracycline (83.3%), which correlates with the antibiotic uses in the farms included in this study [21] as well as follows the global trends of antibiotic resistance in *Streptococcus* spp. [38]. Interestingly, *S. uberis* was sensitive to most of the antibiotics tested in this study with 100% sensitivity to ampicillin, amoxicillin, bacitracin, chloramphenicol, gentamicin, vancomycin; 90% to ceftazidime and 80% to penicillin. On the other hand, *S. agalactiae* showed resistance to penicillin (100%), ceftazidime and gentamicin (66.7%), ampicillin and amoxicillin (33.3%). Increased resistance in *S. agalactiae* is not unusual, and their increased resistance in comparison to *S. uberis* and *S. dysagalactiae* has been reported earlier [5]. Considering the overall resistance pattern, an antibiotic susceptibility testing is suggested before prescribing any antibiotic to treat Streptococcal mastitis.

In this study, the prevalence of *S. uberis* was documented highest (55.6%). Considering the worldwide increasing trend of *S. uberis* in bovine mastitis, this study was further extended to characterize the *S. uberis* isolates through whole genome sequencing (WGS). Sequencing of the five representative isolates followed by assembly revealed the genome sizes ranging from 1886469 bp (strain 2055) to 2033655 bp (strain 2053) with 36.4 to 36.5% GC contents, respectively. The genome sizes correspond to the *S. uberis* genomes reported earlier [28][26]. The variation in genome was investigated and annotation revealed the presence of phages and insertion sequence of variable lengths. The genome of strain 2053 carried two complete phages and a total of around 140 kb phage-related sequences that were absent in the other four genomes, which might have greatly contributed to its increased genome size. Pan-genome analysis, whole genome phylogeny and MLST revealed the presence of two unknown sequence types (STs) of *S. uberis* in the study populations. Similarly, novel ST types have been reported earlier from different countries [28][5][39][26], which might be related to the limited databases of *S. uberis* genomes. The clustering of four out of five BD isolates with India, Thailand and China indicated the circulation of similar STs in this subcontinent.

*S. uberis* is known to establish a successful intramammary infection through the exploitation of different virulence factors associated with invasion of mammary epithelium, evasion of host immunity, binding antimicrobial compounds, biofilm formation through the acquisition of amino acids by breaking milk protein casein [5](. Although the virulence determinant in *S. uberis* is not completely understood; several putative virulence determinants have been described previously [28][26][40]. Considering the previous reports and *S. uberis* putative virulence database (suPVDB), a total of sixty-one (61) putative virulence factors were investigated, where forty-one (41) virulence factors were identified in the studied genomes. Among the virulence factors thirty-five (35) virulence factors were common to all, including genes encoding for major virulence factors relating to the adhesion to epithelial cells (*srtA, fbp54/ rqcH* and *fbps*) [41][42], colonization (*lmb*) [43], invasion/ penetration and dissemination (*pauA/ skc*) [44][45], immuno invasion through the synthesis of hyalurinic acid capsule (*hasC*) and delayed accumulation of lymphocytes (*scpA* and C5a peptidase) [46][47], biofilm formation (*cpsBCD*) [48], hemolytic activity (*cylAG* and hemolysis like protein) [28][49] and virulence regulator (*mga/ vru*) which is known to regulate a number of phenotypes such as growth, biofilm formation and resistance to phagocytosis [50][51].

In addition, five (*cps4E/pglC, cpsM, hasA, hasB* and uberolysin encoding gene) were native to the strains 2002, 20021, 2055 and 2066. HasABC is associated with the formation of the hyaluronic acid capsule, which was required for the prevention of opsonization and phagocytosis as well as escaping neutrophil traps [52]; however, some authors claimed they are non-essential or play a minor role in the induction of mastitis by *S. uberis* [26][53] [54]. Similarly, bacteriocin provides competitive benefits to the bacterium for surviving in a complex environment by inhibiting the growth of other bacteria. The genome 2053 carried five different genes associated with bacteriocin production, similar to the other genome in this study except for uberolysin. The absence of uberolysin might not render strain 2053 avirulent but may give additional benefits or fitness to the other isolates carrying the gene; however, further investigations are necessary to establish the associations.

The *vru* gene was reported to regulate a number of virulence genes, and inactivation of this gene was responsible for reduced mammary gland colonization and clinical signs [55]. A four bp deletion at the upstream (−76 to −79) was speculated to play an important role in the regulatory function of *vru* gene and subsequently influence host-pathogen interactions. In this study, all five genomes had a five bp deletion (TATAA) at −75 to −79 position similar to that reported in EF20, a non-virulent *S. uberis* strain reported from the UK [28]. Considering the putative virulence gene makeup, although strain 2053 closely clustered with EF20, they are genetically different, and the deletion at upstream of the *vru* gene might not be critical to the virulence of the isolates. However, as we did not perform phenotypic studies, making any concrete statement from our findings might be misleading. Thus, further phenotypic experiments with isolates having different variations in the *vru* upstream would provide comprehensive information on its association with the regulation of *vru* gene and virulence of *S. uberis* in bovine mastitis.

## Conclusions

This study in a nutshell, gives a snapshot of the *Streptococcus* spp. inhabiting mammary gland of cattle with clinical mastitis in Bangladesh and introduced two new species *S. hyovaginalis* and *S. urinalis* having a possible association with mastitis for the first time in the world. Genomic information of *S. uberis* generated in this study revealed two novel STs and possible virulence makeup that will significantly contribute to the development of new tools for virulence profiling, ST typing and the development of vaccines for the control of mastitis caused by the emerging pathogen. Moreover, the sequence data generated could be used for the exploration of the genomic background behind the evolution of this pathogen and the story behind its enhanced environmental fitness over time.

## Supporting informations

**S1 Table. Distribution of virulence related genes in *Streptococcus uberis* isolated in this study with corresponding locus in the genomes**

## References

1. Cheng WN, Han SG. Bovine mastitis: risk factors, therapeutic strategies, and alternative treatments - A review. Asian-Australasian J Anim Sci. 2020;33: 1699–1713. doi:10.5713/AJAS.20.0156

2. Bradley AJ. Bovine Mastitis: An Evolving Disease. Vet J. 2002;164: 116–128. doi:10.1053/TVJL.2002.0724

3. Egwu GO, Zaria LT, Onyeyili PA, Ambali AG, Adamu SS, Birdling M. Studies on the microbiological flora of caprine mastitis and antibiotic inhibitory concentrations in Nigeria. Small Rumin Res. 1994;14: 233–239. doi:10.1016/0921-4488(94)90046-9

4. Leigh JA. *Streptococcus uberis*: a permanent barrier to the control of bovine mastitis? Vet J. 1999;157: 225–238. doi:10.1053/TVJL.1998.0298

5. Kabelitz T, Aubry E, van Vorst K, Amon T, Fulde M. The role of *Streptococcus* spp. in bovine mastitis. Microorganisms. 2021;9. doi:10.3390/MICROORGANISMS9071497

6. Cojkic A, Cobanovic N, Suvajdzic B, Savic M, Petrujkic B, Dimitrijevic V. Chronic mastitis in cows caused by *Streptococcus dysgalactiae*: Case report. Vet Glas. 2015;69: 303–314. doi:10.2298/VETGL1504303C

7. Jain B, Tewari A, Bhandari BB, Jhala MK. Antibiotic resistance and virulence genes in *Streptococcus agalactiae* isolated from cases of bovine subclinical mastitis. Vet Arh. 2012;82: 423–432.

8. Nikolova M, Urumova V, Liuzkanov M. *Streptococcus* spp. as etiological agent of subclinical and clinical mastitis of dairy cows in Republic of Bulgaria. Trakia J Sci 2022;20: 113–118. doi:10.15547/TJS.2022.02.005

9. Alnakip MEA, Rhouma NR, Abd-Elfatah EN, Quintela-Baluja M, Böhme K, Fernández-No I, et al. Discrimination of major and minor streptococci incriminated in bovine mastitis by MALDI-TOF MS fingerprinting and 16S rRNA gene sequencing. Res Vet Sci. 2020;132: 426–438. doi:10.1016/J.RVSC.2020.07.027

10. Carvalho-Castro GA, Silva JR, Paiva L V., Custódio DAC, Moreira RO, Mian GF, et al. Molecular epidemiology of *Streptococcus agalactiae* isolated from mastitis in Brazilian dairy herds. Braz J Microbiol. 2017;48: 551–559. doi:10.1016/J.BJM.2017.02.004

11. Benić M, Maćešić N, Cvetnić L, Habrun B, Cvetnić Ž, Turk R, et al. Bovine mastitis: a persistent and evolving problem requiring novel approaches for its control – a review. Vet Arh. 2018;88: 535–557. doi:10.24099/VET.ARHIV.0116

12. Hillerton JE, Berry EA. The management and treatment of environmental streptococcal mastitis. Vet Clin North Am Food Anim Pract. 2003;19: 157–169. doi:10.1016/S0749-0720(02)00069-5

13. Erskine RJ, Wagner S, DeGraves FJ. Mastitis therapy and pharmacology. Vet Clin North Am Food Anim Pract. 2003;19: 109–138. doi:10.1016/S0749-0720(02)00067-1

14. Kalińska A, Gołębiewski M, Wójcik A. Mastitis pathogens in dairy cattle - a review. World Sci News. 2017; 22–31.

15. Cvetnic L, Samardžija M, Habrun B, Kompes G, Benić M. Microbiological monitoring of mastitis pathogens in the control of udder health in dairy cows. 2016;53: 131–140.

16. Phuektes P, Mansell PD, Dyson RS, Hopper ND, Dick JS, Browning GF. Molecular epidemiology of *Streptococcus uberis* isolates from dairy cows with mastitis. J Clin Microbiol. 2001;39: 1460–1466. doi:10.1128/JCM.39.4.1460-1466.2001

17. Hasan MT, Islam MR, Runa NS, Hasan MN, Uddin AHMM, Singh SK. Study on bovine sub-clinical mastitis on farm condition with special emphasis on antibiogram of the causative bacteria. Bangladesh J Vet Med. 2017;14: 161–166. doi:10.3329/BJVM.V14I2.31386

18. Rahman MM, Islam MR, Uddin MB, Aktaruzzaman M. Prevalence of subclinical mastitis in dairy cows reared in Sylhet district of Bangladesh. Int J BioResearch. 2010;1: 23–28.

19. Rahman MT, Islam MS, Hasan M. Isolation and identification of bacterial agents causing clinical mastitis in cattle in Mymensingh and their antibiogram profile. Microbes Heal. 2013;2: 19–21. doi:0.3329/mh.v2i1.17258

20. Bag MAS, Arif M, Riaz S, Khan MSR, Islam MS, Punom SA, et al. Antimicrobial resistance, virulence profiles, and public health significance of *Enterococcus faecalis* isolated from clinical mastitis of cattle in Bangladesh. BiomedRes Int. 2022;2022. doi:10.1155/2022/8101866

21. Bag MAS, Khan MSR, Sami MDH, Begum F, Islam MS, Rahman MM, et al. Virulence determinants and antimicrobial resistance of *E. coli* isolated from bovine clinical mastitis in some selected dairy farms of Bangladesh. Saudi J Biol Sci. 2021;28: 6317–6323. doi:10.1016/J.SJBS.2021.06.099

22. Hahne J, Kloster T, Rathmann S, Weber M, Lipski A. Isolation and characterization of *Corynebacterium* spp. from bulk tank raw cow’s milk of different dairy farms in Germany. PLoS One. 2018;13. doi:10.1371/JOURNAL.PONE.0194365

23. Bauer AW, Kirby WM, Sherris JC, Turch M. Antibiotic susceptibility testing by a standardized single disk method. Am J Clin Pathol. 1966;45: 493–496.

24. CLSI M100-Ed30 - Performance Standards for Antimicrobial Susceptibility Testing - 30th Edition. [cited 3 Oct 2022]. Available: https://webstore.ansi.org/Standards/CLSI/CLSIM100Ed30

25. Magiorakos AP, Srinivasan A, Carey RB, Carmeli Y, Falagas ME, Giske CG, et al. Multidrug-resistant, extensively drug-resistant and pandrug-resistant bacteria: an international expert proposal for interim standard definitions for acquired resistance. Clin Microbiol Infect. 2012;18: 268–281. doi:10.1111/J.1469-0691.2011.03570.X

26. Vezina B, Al-harbi H, Ramay HR, Soust M, Moore RJ, Olchowy TWJ, et al. Sequence characterisation and novel insights into bovine mastitis-associated *Streptococcus uberis* in dairy herds. Sci Rep. 2021;11. doi:10.1038/S41598-021-82357-3

27. Alcantara RB, Read RDA, Valderas MW, Brown TD, Roop RM. Intact purine biosynthesis pathways are required for wild-type virulence of *Brucella abortus* 2308 in the BALB/c mouse model. Infect Immun. 2004;72: 4911–4917. doi:10.1128/IAI.72.8.4911-4917.2004

28. Hossain M, Egan SA, Coffey T, Ward PN, Wilson R, Leigh JA, et al. Virulence related sequences; insights provided by comparative genomics of *Streptococcus uberis* of differing virulence. BMC Genomics. 2015;16. doi:10.1186/S12864-015-1512-6

29. Kaczorek E, Małaczewska J, Wójcik R, Siwicki AK. Biofilm production and other virulence factors in *Streptococcus* spp. isolated from clinical cases of bovine mastitis in Poland. BMC Vet Res. 2017;13. doi:10.1186/S12917-017-1322-Y

30. Contreras-Moreira B, Vinuesa P. GET_HOMOLOGUES, a versatile software package for scalable and robust microbial pangenome analysis. Appl Environ Microbiol. 2013;79: 7696–701. doi:10.1128/AEM.02411-13

31. Page AJ, Cummins CA, Hunt M, Wong VK, Reuter S, Holden MTG, et al. Roary: rapid large-scale prokaryote pan genome analysis. Bioinformatics. 2015;31: 3691–3693. doi:10.1093/BIOINFORMATICS/BTV421

32. Bradley AJ, Leach KA, Breen JE, Green LE, Green MJ. Survey of the incidence and aetiology of mastitis on dairy farms in England and Wales. Vet Rec. 2007;160: 253–258. doi:10.1136/VR.160.8.253

33. Petrovski KR, Heuer C, Parkinson TJ, Williamson NB. The incidence and aetiology of clinical bovine mastitis on 14 farms in Northland, New Zealand. N Z Vet J. 2009;57: 109–115. doi:10.1080/00480169.2009.36887

34. Verbeke J, Piepers S, Supré K, De Vliegher S. Pathogen-specific incidence rate of clinical mastitis in Flemish dairy herds, severity, and association with herd hygiene. J Dairy Sci. 2014;97: 6926–6934. doi:10.3168/jds.2014-8173

35. Zhang H, Yang F, Li X pu, Luo J yin, Wang L, Zhou Y long, et al. Detection of antimicrobial resistance and virulence-related genes in *Streptococcus uberis* and *Streptococcus parauberis* isolated from clinical bovine mastitis cases in northwestern China. J Integr Agric. 2020;19: 2784–2791. doi:10.1016/S2095-3119(20)63185-9

36. Gomes F, Henriques M. Control of Bovine Mastitis: Old and Recent Therapeutic Approaches. Curr Microbiol. 2016;72: 377–382. doi:10.1007/S00284-015-0958-8

37. Bonsaglia ECR, Gomes MS, Canisso IF, Zhou Z, Lima SF, Rall VLM, et al. Milk microbiome and bacterial load following dry cow therapy without antibiotics in dairy cows with healthy mammary gland. Sci Rep. 2017;7. doi:10.1038/S41598-017-08790-5

38. Haenni M, Lupo A, Madec J-Y. Antimicrobial resistance in *Streptococcus* spp. Microbiol Spectr. 2018;6. doi:10.1128/MICROBIOLSPEC.ARBA-0008-2017

39. Silva NCC, Yang Y, Rodrigues MX, Tomazi T, Bicalho RC. Whole-genome sequencing reveals high genetic diversity of *Streptococcus uberis* isolated from cows with mastitis. BMC Vet Res. 2021;17. doi:10.1186/S12917-021-03031-4

40. Zouharova M, Nedbalcova K, Slama P, Bzdil J, Masarikova M, Matiasovic J. Occurrence of virulence-associated genes in *Streptococcus uberis* and *Streptococcus parauberis* isolated from bovine mastitis. Vet Med (Praha). 2022;67 (2022): 123–130. doi:10.17221/95/2021-VETMED

41. Lalioui L, Pellegrini E, Dramsi S, Baptista M, Bourgeois N, Doucet-Populaire F, et al. The SrtA Sortase of *Streptococcus agalactiae* is required for cell wall anchoring of proteins containing the LPXTG motif, for adhesion to epithelial cells, and for colonization of the mouse intestine. Infect Immun. 2005;73: 3342–3350. doi:10.1128/IAI.73.6.3342-3350.2005

42. Terao Y, Kawabata S, Kunitomo E, Murakami J, Nakagawa I, Hamada S. Fba, a novel fibronectin-binding protein from *Streptococcus pyogenes*, promotes bacterial entry into epithelial cells, and the fba gene is positively transcribed under the Mga regulator. Mol Microbiol. 2001;42: 75–86. doi:10.1046/J.1365-2958.2001.02579.X

43. Spellerberg B, Rozdzinski E, Martin S, Weber-Heynemann J, Schnitzler N, Lütticken R, et al. Lmb, a protein with similarities to the LraI adhesin family, mediates attachment of *Streptococcus agalactiae* to human laminin. Infect Immun. 1999;67: 871–878. doi:10.1128/IAI.67.2.871-878.1999

44. Leigh JA. Activation of bovine plasminogen by *Streptococcus uberis*. FEMS Microbiol Lett. 1993;114: 67–71. doi:10.1111/J.1574-6968.1993.TB06552.X

45. Ward PN, Field TR, Rapier CD, Leigh JA. The activation of bovine plasminogen by PauA is not required for virulence of *Streptococcus uberis*. Infect Immun. 2003;71: 7193–7196. doi:10.1128/IAI.71.12.7193-7196.2003/ASSET/56F2D591-2662-4867-8FE3-432A85E43C39/ASSETS/GRAPHIC/II1231088002.JPEG

46. O’Connor SP, Patrick Cleary P. In vivo *Streptococcus pyogenes* C5a peptidase activity: analysis using transposon- and nitrosoguanidine-induced mutants. J Infect Dis. 1987;156: 495–504. doi:10.1093/INFDIS/156.3.495

47. Ward PN, Field TR, Ditcham WGF, Maguin E, Leigh JA. Identification and disruption of two discrete loci encoding hyaluronic acid capsule biosynthesis genes *hasA*, *hasB*, and *hasC* in *Streptococcus uberis*. Infect Immun. 2001;69: 392–399. doi:10.1128/IAI.69.1.392-399.2001

48. Morona JK, Miller DC, Morona R, Paton JC. The effect that mutations in the conserved capsular polysaccharide biosynthesis genes *cpsA*, *cpsB*, and *cpsD* have on virulence of *Streptococcus pneumoniae*. J Infect Dis. 2004;189: 1905–1913. doi:10.1086/383352

49. Spellerberg B, Martin S, Brandt C, LÃ¼tticken R. The *cyl* genes of *Streptococcus agalactiae* are involved in the production of pigment. FEMS Microbiol Lett. 2000;188: 125–128. doi:10.1111/J.1574-6968.2000.TB09182.X

50. Cho KH, Caparon MG. Patterns of virulence gene expression differ between biofilm and tissue communities of *Streptococcus pyogenes*. Mol Microbiol. 2005;57: 1545–1556. doi:10.1111/J.1365-2958.2005.04786.X

51. Le Breton Y, Belew AT, Freiberg JA, Sundar GS, Islam E, Lieberman J, et al. Genomewide discovery of novel M1T1 group A streptococcal determinants important for fitness and virulence during soft-tissue infection. PLoS Pathog. 2017;13. doi:10.1371/JOURNAL.PPAT.1006584

52. Cole JN, Pence MA, von Köckritz-Blickwede M, Hollands A, Gallo RL, Walker MJ, et al. M protein and hyaluronic acid capsule are essential for *in vivo* selection of *covRS* mutations characteristic of invasive serotype M1T1 group A Streptococcus. MBio. 2010;1. doi:10.1128/MBIO.00191-10

53. Field TR, Ward PN, Pedersen LH, Leigh JA. The hyaluronic acid capsule of *Streptococcus uberis* is not required for the development of infection and clinical mastitis. Infect Immun. 2003;71: 132–139. doi:10.1128/IAI.71.1.132-139.2003

54. Ward PN, Holden MTG, Leigh JA, Lennard N, Bignell A, Barron A, et al. Evidence for niche adaptation in the genome of the bovine pathogen *Streptococcus uberis*. BMC Genomics. 2009;10. doi:10.1186/1471-2164-10-54

55. Flores AR, Olsen RJ, Wunsche A, Kumaraswami M, Shelburne SA, Carroll RK, et al. Natural variation in the promoter of the gene encoding the Mga regulator alters host-pathogen interactions in group a Streptococcus carrier strains. Infect Immun. 2013;81: 4128–4138. doi:10.1128/IAI.00405-13

